# Computational characteristics of the striatal dopamine system described by reinforcement learning with fast generalization

**DOI:** 10.1101/2019.12.12.873950

**Authors:** Yoshihisa Fujita, Sho Yagishita, Haruo Kasai, Shin Ishii

## Abstract

Generalization enables applying past experience to similar but nonidentical situations. Therefore, it may be essential for adaptive behaviors. Recent neurobiological observation indicates that the striatal dopamine system achieves generalization and subsequent discrimination by updating corticostriatal synaptic connections in differential response to reward and punishment. To analyze how the computational characteristics in this system affect behaviors, we proposed a novel reinforcement learning model with multilayer neural networks in which the synaptic weights of only the last layer are updated according to the prediction error. We set fixed connections between the input and hidden layers so as to maintain the similarity of inputs in the hidden-layer representation. This network enabled fast generalization, and thereby facilitated safe and efficient exploration in reinforcement learning tasks, compared to algorithms which do not show generalization. However, disturbance in the network induced aberrant valuation. In conclusion, the unique computation suggested by corticostriatal plasticity has the advantage of providing safe and quick adaptations to unknown environments, but on the other hand has the potential defect which can induce maladaptive behaviors like delusional symptoms of psychiatric disorders.

**Author summary:** The brain has an ability to generalize knowledge obtained from reward- and punishment-related learning. Animals that have been trained to associate a stimulus with subsequent reward or punishment respond not only to the same stimulus but also to resembling stimuli. How does generalization affect behaviors in situations where individuals are required to adapt to unknown environments? It may enable efficient learning and promote adaptive behaviors, but inappropriate generalization may disrupt behaviors by associating reward or punishment with irrelevant stimuli. The effect of generalization here should depend on computational characteristics of underlying biological basis in the brain, namely, the striatal dopamine system. In this research, we made a novel computational model based on the characteristics of the striatal dopamine system. Our model enabled fast generalization and showed its advantage of providing safe and quick adaptation to unknown environments. By contrast, disturbance of our model induced abnormal behaviors. The results suggested the advantage and the shortcoming of generalization by the striatal dopamine system.

## Introduction

Animals’ survival with reward-seeking behaviors should be accompanied with risks. Outcome observation resulting from the pairing of a current state and a taken action could be a clue to ensure optimal behaviors, but it may be associated with substantial energy consumption and aversive experience. Such a learning process is inefficient and even harmful, especially when animals are required to adapt to new environments. Animals instead generalize their past experience for predicting outcome, even in novel situations. Generalization may be essential for efficient adaptive behaviors, whereas generalization to an abnormal extent can be maladaptive and is implicated in psychiatric disorders [1–3].

Psychological studies have investigated generalization by using the conditioning paradigm in behavioral experiments [4]. If a response has been established by a stimulus paired with an outcome (i.e., reward or punishment), then resembling stimuli evoke similar responses. This “law of effect” depends on the extent that the second stimulus resembles the first stimulus, and is termed “stimulus generalization” [4, 5]. Discrimination can then occur if the first stimulus is paired with a reward but the resembling stimulus is paired with no reward; as a consequence, only the first stimulus elicits a response. How the brain establishes stimulus generalization has been explained, based on artificial neural networks [6–9]. Previous studies [6–9] have focused on the stimulus–response relationship, although generalization can be incorporated in the interaction between individuals and the environment. Reinforcement learning involves this interaction and is used as a model of reward-driven learning [10–12]. In the field of artificial intelligence, reinforcement learning in combination with artificial neural networks achieves a high performance, which suggests the contribution of generalization to adaptive behaviors [13]. However, neurobiological evidence indicates that the neural system has a unique computation for reward-driven learning and generalization, compared to algorithms used for artificial intelligence [14]. This fact raises the question regarding how its computational characteristics differ from those of ordinary algorithms. Advantages and shortcomings should exist with regard to adaptive behaviors.

Theoretical attempts have focused on separate systems that process positive and negative values in the brain [15–18], which is not adopted when using ordinary reinforcement learning. Dopamine and its main target, the striatum, have central roles in reward- and punishment-related learning [19]. Dopamine neurons show a positive response to a greater-than-expected reward and a negative response to a less-than-expected reward, which indicates that dopamine codes reward prediction error [20]. The striatum receives dopamine signals and glutamatergic input from the cortex and thalamus [21]. Dopamine modulates synaptic plasticity between the cortex and the striatum during reward-related learning [22]. The striatum is primarily composed of spiny projection neurons (SPNs), which can be divided into SPNs that primarily express the dopamine D1 receptor (D1-SPNs) and SPNs that express the dopamine D2 receptor (D2-SPNs) [23]. The D1-SPNs respond to the phasic increase in dopamine [24], whereas D2-SPNs may respond to the phasic decrease in dopamine [25]. Perturbing the activity of the D1- and D2-SPNs inhibits reward learning and punishment learning, respectively [25]. The advantages of having such separate systems have been discussed in computational studies [26, 27]. One study [26] demonstrated an advantage in learning reward uncertainty. Another study [27] showed the possibility of achieving safe behaviors.

However, recent neural recordings and optogenetic manipulations provide some data that suggests different roles of SPNs from those in existing models [28]. Our new experiments with a classical conditional task found that D1- and D2-SPNs are differentially responsible for stimulus generalization and discrimination (Yagishita, Kasai and others, unpublished). The same series of experiments revealed that stimulus generalization/discrimination occurred solely by dopamine-dependent plasticity of SPN spines that receive cortical inputs, which implies that updating in other connections (e.g., intracortical synaptic connections) would be minor. These observations suggest learning rules that update the synaptic weights of the last layer in a multilayer neural network—in our case, corticostriatal connections—are essential. Such learning rules actually enable remarkably fast learning in reservoir computing (RC) and extreme learning machine (ELM) [29–31]. Reservoir computing and ELM are neural networks that train only their readouts (i.e., the last layer connections); RC has recurrent connections, whereas ELM does not.

In this study, we propose a novel reinforcement learning model that reproduces stimulus generalization and discrimination, while accounting for the physiological characteristics of striatal SPNs. We used a neural network for value estimation and introduced fixed connections between the input and hidden layers, as in RC and ELM. Reservoir computing has the potential for context-dependent value estimation because of its recurrent connections. However, for simplicity, we adopted a feed-forward neural network with the same architecture as ELM. The extent of stimulus generalization depends on the similarity of stimuli; therefore, we did not use random and fixed connections implemented in ordinary ELM. We instead set the fixed connections so as to maintain the similarity of inputs in the hidden-layer representation. In addition, we introduced connections, the weights of which are separately updated by positive and negative reward prediction errors, while taking into account the differential roles of D1- and D2-SPNs in the striatum. This neural network enabled generalization in a quick manner. We named this model “Outspread Valuation for Reward Learning and Punishment Learning” (“OVaRLAP”). The OVaRLAP model achieved safe and efficient exploration in painful grid-world navigation tasks. This achievement suggested that quick generalization contributed to safe and efficient reward-seeking and pain-avoiding. We also tested disturbed OVaRLAP to examine whether abnormal generalization underlies maladaptive behaviors, as implied in psychological and psychiatric studies [3, 32, 33]. The OVaRLAP method with noisy generalization and impaired discrimination induced aberrant valuation, after repeating reward-seeking behaviors. These results showed that the unique computation suggested by the striatal dopamine system enabled safe and efficient exploration, but on the other hand had the potential defect which can cause maladaptive behaviors.

## Methods

We developed OVaRLAP to analyze how the computational characteristics in the striatal dopamine system affect behaviors. We first evaluated OVaRLAP in a spatial navigation task in painful grid-worlds by comparing its performance with those of two other representative algorithms (Fig 1A-1B). We then examined behaviors of disturbed OVaRLAP in a spatial navigation task in a painless grid-world (Fig 1C-1D).

**Fig 1.**
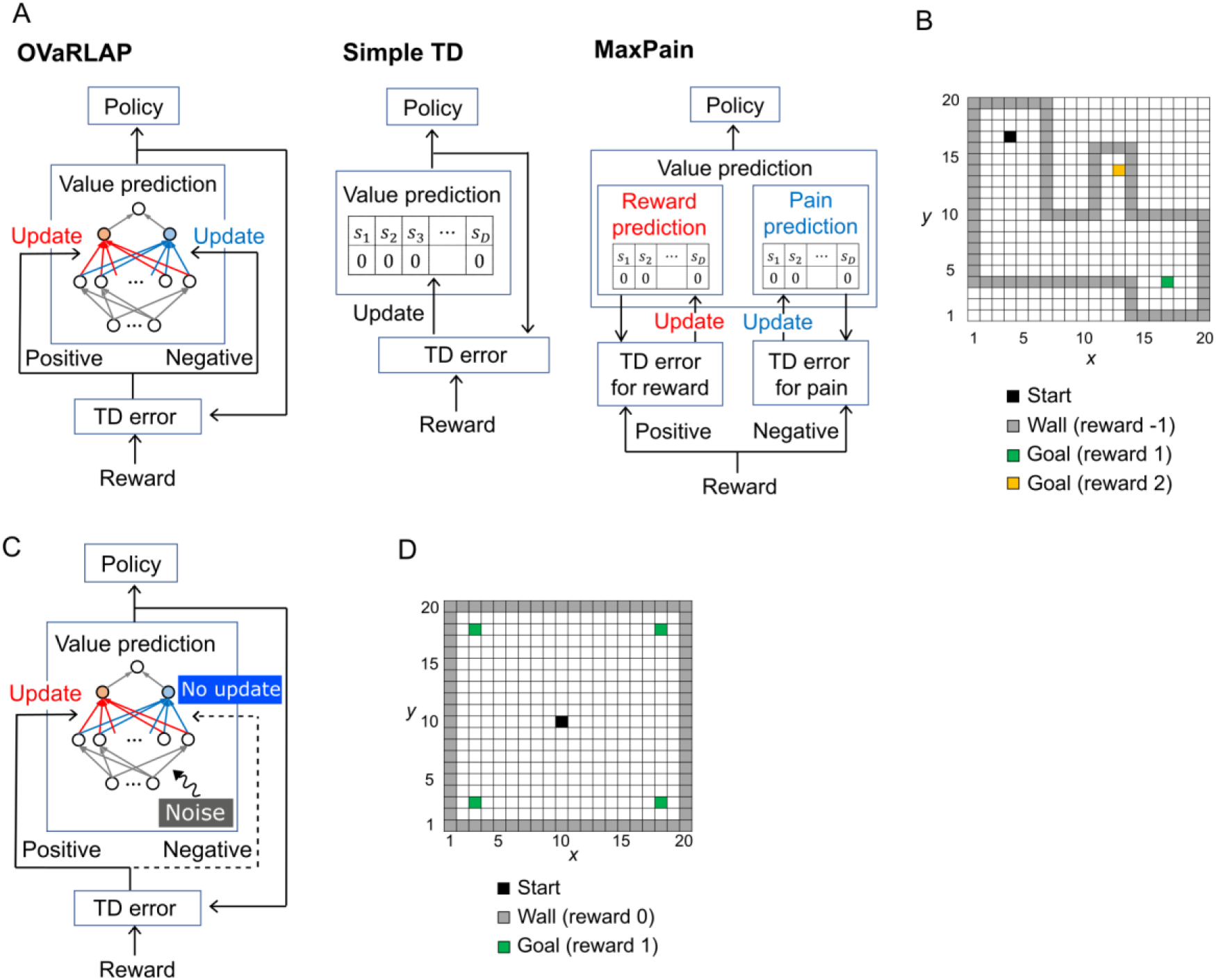
The architecture of TD learning algorithms and the task. (**A**) Schematic diagrams of OVaRLAP (left), simple TD learning (middle), and MaxPain (right). In OVaRLAP, the value prediction is conducted using a neural network with fixed connections (gray arrows) and distinct connections updated by positive and negative TD errors (red and blue arrows, respectively). In our implementations of simple TD learning and MaxPain, the predicted value is represented by lookup tables in which the state, *s_k_*, indicates a position (*x*, *y*) in the two-dimensional grid-world. (**B**) The navigation task in the two-dimensional grid-world. The nongray squares and gray squares indicate passable and not passable, respectively. The black square is the starting position, and the blue and yellow squares are the goals at which the agent receives the positive rewards of 1 and 2, respectively. When the agent receives a positive reward, an episode ends so that the agent restarts the task from the starting position. If an agent hits a wall (i.e., a not passable square), it receives a negative reward of −1, but continues the task by staying at the same square. (**C**) Schematic diagram of OVaRLAP in which updating connections, based on the negative TD error (blue connections), are impaired. In addition, noise was introduced to induce an anomaly in the initialization of the fixed connections between the input and the hidden layers (for the details of noise here, see the Methods section). (**D**) The navigation task in the two-dimensional grid-world. The black square is the starting position. The green squares are the goals at which the agent receives a positive reward of 1. If the agent receives a positive reward, a single episode ends, and a new episode restarts from the starting position. In this task, when the agent hits a wall (i.e., a gray square), no negative reward is given, and the agent remains at the same square.

### Model description of OVaRLAP

A neural network was used for value estimation in OVaRLAP. It consists of an input layer, a hidden layer, two pre-output neurons, and an output neuron (Fig 1A, left). The input represents the state at discretized time, *t*. We applied our learning method to two types of navigation tasks, both in two-dimensional grid-worlds, in which position (*x*, *y*), *x* ∈ {1, …, 20}, and *y* ∈ {1, … 20} were simply represented by a two-dimensional index function:

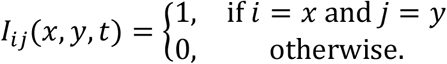

The input signals were transformed by fixed connections into hidden-layer activity patterns. We set the number of hidden-layer neurons and the weight of fixed connections so as to reproduce the shape of the generalization gradient, which has been commonly observed across various species, behavioral contexts, and sensory modalities [4]. The number of hidden-layer neurons was set to 900. The hidden-layer activity *h_k_*(*x*, *y*, *t*) for *k* = 1, …, 900 was given by

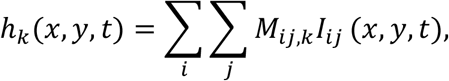

in which *M*_*ij*,*k*_ is the weight of the fixed connection from the input-layer neuron (*i*, *j*) to the hidden-layer neuron *k*. Based on the definition of *I*_*ij*_(*x*, *y*, *t*), *h*_*k*_(*x*, *y*, *t*) was also given by

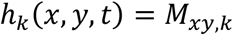

in which (*x*, *y*) is the agent’s position at time *t*.

For the generalization capability, we assumed the hidden-layer representation becomes similar when the input signal is similar. In our case, the values of *h_k_*(*x*_1_, *y*_1_, *t*) and *h_k_*(*x*_2_, *y*_2_, *t*) are similar if position (*x*_1_, *y*_1_) is close to position (*x*_2_, *y*_2_), based on Euclidian distance. To reflect this request, we set the fixed connections *M*_*ij*,*k*_ to follow the two-dimensional Gaussian function:

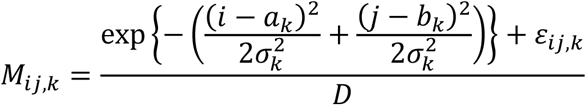

in which *D* is the number of the input-layer neurons used for normalization (*D* = 400, because of the 20 × 20 grid-worlds); *a*_*k*_ and *b*_*k*_ denote the center and are set as *a*_*k*_ ∈ {1, …, 20}, *b*_*k*_ ∈ {1, …, 20}, and 20(*a*_*k*_ − 1) + *b*_*k*_ = *ceil*(400*k*/900) so that the center location was linearly correlated in the 2D space; 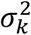 is the variance sampled from a log-normal distribution, 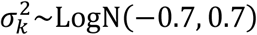; and *ε*_*ij*,*k*_ is the noise, as defined later. The distribution of 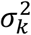 resulted that the hidden-layer neurons ranged from neurons that responded to specific input to neurons that responded to a wide range of input. This setting is consistent with physiological observations [34]. The noise *ε*_*ij*,*k*_ is unnecessary in the normal case; however, we introduced it for the purpose of analyzing the effect of abnormal perturbation in OVaRLAP (Fig 1C). When simulating abnormal perturbation, the noise ε_*ij*,*k*_ was added by using the following formulas:

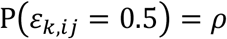

and

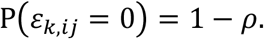

No noise apparently exists if *ρ* is zero. This noise was initially introduced but not changed through the learning process. In our simulation experiment, we applied different patterns of noise *ε*_*ij*,*k*_ under a specific value of *ρ*, and examined the collective behaviors of the learning.

The two pre-output neurons then received signals from the hidden-layer neurons. For *m* = 1,2, the activities of the pre-output neurons *d*_*m*_(*x*, *y*, *t*) are given by

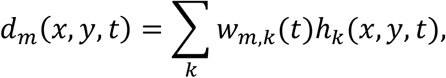

in which *W_m,k_*(*t*) is the weight of the connection from hidden-layer neuron *k* to the pre-output neuron *m*. In this instance, *d*_1_(*x*, *y*, *t*) and *d*_2_(*x*, *y*, *t*) represented the positive and negative values, respectively. The final output of the network *v*(*x*, *y*, *t*) integrated these values, as follows:

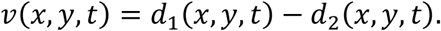

During the interaction between the agent and the environment, the connection weights *W_m,k_*(*t*) were updated, depending on the prediction error, *δ*. To calculate *δ*, we used an action-value function *Q*(*s*, *a*, *t*) for state *s* and action *a* at time *t* [35]. We defined *Q*(*s*, *a*, *t*) by using the output of the network, as follows:

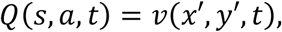

in which *s* = (*x*, *y*), *a* = (*Δx*, *Δy*), *x*′ = *x* + *Δx*, and *y*′, = *y* + *Δy*. In our navigation tasks, (*Δx*, *Δy*) = (1,0), (−1,0), (0, −1), or (0,1), if the action was effective. The prediction error *δ* was given by

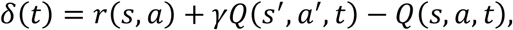

in which *r*(*s*, *a*) is the actual reward, *s*′ denotes the state at time *t* + 1, *a*′ denotes the action at time *t* + 1, and *γ* is the discount factor. We used temporal difference (TD) learning for the action-value function (i.e., state–action–reward–state–action [SARSA]) [36] because the agent needed to remain at the same square after an ineffective action (i.e., hitting a wall). If we used a simple TD for the state-value function, such ineffective actions would introduce noise to the value learning. The connection weight *W*_1,*k*_(*t*) was updated only when *δ*(*t*) was positive, and *W*_2,*k*_(*t*) was updated only when *δ*(*t*) was negative. In the actual implementation, the updating rules were as follows:

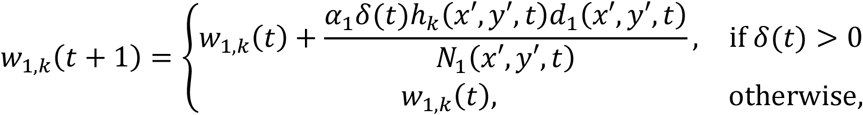

and

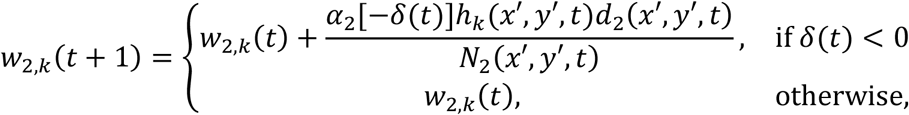

in which *α*_1_ and *α*_2_ are the learning rates and *N*_1_(*x*, *y*, *t*) and *N*_2_(*x*, *y*, *t*) are the normalization terms, which are given by

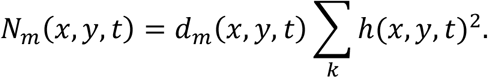

for *m* = 1,2. These normalization terms are introduced to make sure that

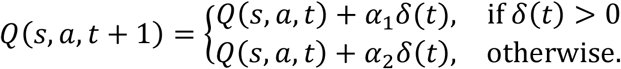

### Algorithms for comparison

We also implemented two representative algorithms to compare with the OVaRLAP model. One algorithm was simple TD learning [35] (Fig 1A, middle); in our particular case, the algorithm was SARSA. In our implementation, a value function of states, *v*_*s*_(*x*, *y*, *t*), was represented as a look-up table and updated as follows:

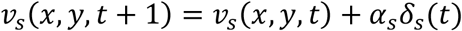

in which *α*_*s*_ is the learning rate and *δ*_*s*_ is the prediction error, called the “TD error.” This error is given by

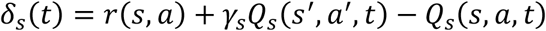

in which *γ*_*s*_ is the discount factor. The action-value function, *Q_*s*_*(*s*, *a*, *t*), is given by

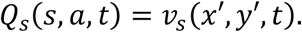

The other algorithm is the MaxPain algorithm [27] (Fig 1A, right). This method is characterized by its distinct systems for the learning values for reward and pain (or punishment). The policy is dependent on the linear combination of the two value functions. In our implementation, we slightly modified the originally proposed MaxPain algorithm to make it comparable with the other methods, while maintaining the essential idea of MaxPain.

First, we used state-value functions of states *v_r_*(*x*, *y*, *t*) and *v_p_*(*x*, *y*, *t*), and their linear combination *v_L_*(*x*, *y*, *t*), to define the action-value functions *Q_r_*(*s*′, *a*′, *t*), *Q_p_*(*s*′, *a*′, *t*), and *Q_L_*(*s*′, *a*′, *t*), as follows:

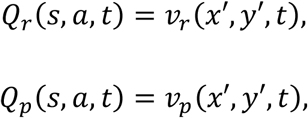

and

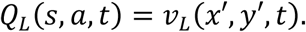

Second, we set the linear combination *v_L_*(*x*, *y*, *t*) without normalization, as follows:

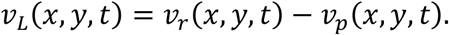

The couple of state-value functions were implemented as look-up tables and updated as

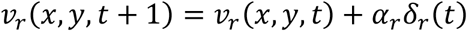

and

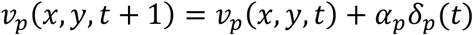

in which *α_r_* and *α_p_* are the learning rates for reward and pain, respectively, and *δ_r_* and *δ_p_* are the prediction errors for reward and pain, respectively. The prediction errors were calculated, depending on whether the reward observation was positive or negative, as follows:

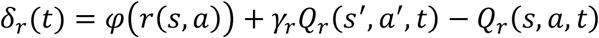

and

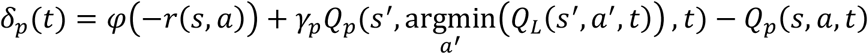

in which *φ*(*z*) = max (*z*, 0) and *γ_r_* and *γ_p_* are the discount factors. These update rules were the same as those in the original study. The pain prediction error, *δ_p_*, was used for maximizing future pain (i.e., for predicting the worst case).

### Action selection

We used the softmax behavioral policy consistently for the three methods. It depends on the value function 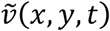 [i.e., *v*(*x*, *y*, *t*) for OVaRLAP, *v_s_*(*x*, *y*, *t*) for the simple TD learning, and *v_L_*(*x*, *y*, *t*) for MaxPain]. The probability that the agent selects an action *a* at position (*x*, *y*) at time *t* is given by

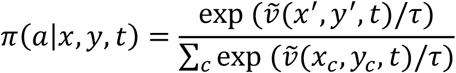

in which *x*′ and *y*′ denote the new state after selecting action *a*, *x_c_* and *y_c_* denote the new state after selecting one of the possible actions, and *τ* is the temperature that controls the trade-off between exploration and exploitation. In our implementation, we used the common *τ* = 0.5, for the three learning methods.

### Painful grid-world navigation task

The purpose of this task was to navigate from the starting position to either of the two goals, while avoiding hitting the wall. Possible actions at each time step were moving one step north, south, east, and west. For example, if the agent moved one step north, (*Δx*, *Δy*) = (0,1). Two goals exist with reward of 1 or 2. If the agent hits a wall, then it received a negative reward of −1 and remained at the same position. This was an exceptional case. In other cases, the agent could by necessity move to the next square. An episode began when the agent started from the starting position and ended when the agent reached either of the goals. The agent repeated such learning episodes.

A single run consisted of 500 learning episodes, after initializing the value function so that the value for each state was zero. We ran 50 separate runs each for OVaRLAP, simple TD learning, and MaxPain with a single grid-world configuration. We conducted five simulation experiments for each of which we used a grid-world with a consistent character but different configuration. The structure of each grid-world is shown in Fig 1B and in S1 Fig. We set the metaparameters for each algorithm, as follows: ρ = 0, *α*_1_ = *α*_2_ = 0.1, and *γ* = 0.95 for OVaRLAP; *α_s_* = 0.1 and *γ_s_* = 0.95 for simple TD learning; and *α_r_* = *α_p_* = 0.1, *γ_r_* = 0.95, and *γ_p_* = 0.5 for MaxPain. OVaRLAP and MaxPain are extended algorithms, based on simple TD; therefore, they use common metaparameters with simple TD, when they could share them.

The five configurations in S1 Figure were generated, based on the following rules, while avoiding symmetric or similar structures. Each configuration had a wide passage from which a narrow passage branched off. These passages had fixed widths and lengths. The starting position and the goal with a reward of 1 were at either end of a wide passage. A goal with reward of 2 was at the end of a narrow passage. S1 Fig. legend has more details of the rules.

### Grid-world navigation task for disturbed OVaRLAP

The purpose of this task was to navigate from the starting position to either of four goals. The structure of the grid-world is shown in Fig 1D. Possible actions at each time step were moving one step north, south, east, and west. Multiple goals existed, each of which gave a reward of 1. The topic in this study was not on risk aversion; therefore, we did not apply a negative reward to the actions that resulted in hitting a wall (i.e., no pain). The agent simply remained at the same position after hitting a wall. Even in this no pain setting, the agents after reinforcement learning likely avoid wall hits, due to the discount factor in the value functions. An episode began when the agent moved from the starting position and ended when the agent reached one of the four goals. A single run consisted of a total of 40,000 time steps (i.e., 378.8 learning episodes on average) after the value function initialization at the onset of the first learning episode. We ran 50 separate runs for OVaRLAP for each of the following settings: normal or no update for a negative TD error (i.e., “intact” or “impaired”) and with or without noise in the fixed connections (i.e., “noised” or “unnoised”). We set the metaparameters, as follows: *ρ* = 0.005 for the noised case and *ρ* = 0 for the unnoised case; *α_1_* = 0.1; *α_2_* = 0.1 for the intact setting (which was the same as in the previous experiment) and *α_2_* = 0 for the impaired setting; *γ* = 0.8; and *τ* = 0.5. In the noised setting, the noise, *ε_ij,k_*, was generated independently for each run and was fixed within each run.

## Results

### The behavior of OVaRLAP

We evaluated OVaRLAP in a spatial navigation task in painful grid-worlds by comparing its performance with those of simple TD learning and MaxPain [27]. These algorithms are schematically shown in Fig 1A. On account that OVaRLAP utilizes a fixed neural network (gray arrows in Fig 1A, left) for the preprocessing of the value learning network (blue and red arrows in Fig 1A, left), it achieves stimulus generalization, based on the similarity of states (Fig 2, left). By contrast, the other two algorithms, which have look-up tables, show no stimulus generalization (Fig 2, middle and right). The task in this study was to seek rewarded goals while avoiding painful wall hits in which the agent had to manage the trade-off between obtaining a reward and avoiding pain (Fig 1B). We used five different grid-worlds, each of which had a safe but lowly rewarded goal and a risky but highly rewarded goal (S1 Fig).

**Fig 2.**
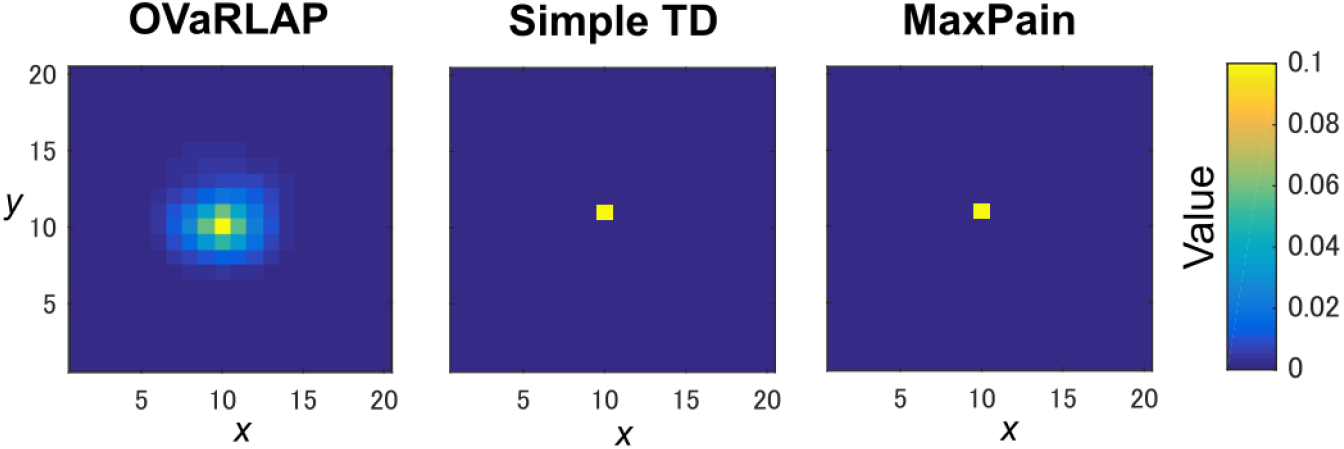
State values after a single update. The value function is updated after receiving a positive reward of 1 once with OVaRLAP (left), simple TD learning (middle), and MaxPain (right) at the center of a two-dimensional grid-world in Fig 1B.

Figure 3A-3C show the average learning curves at each episode over 50 independent runs for the grid-world shown in Fig 1B. The reward per step of the OVaRLAP agent was higher than that of the MaxPain agent, but it did not exceed that of the simple TD learning agent (Fig 3A). Compared to the other agents, the OVaRLAP agent reached each of the goals with a smaller number of steps in the early learning phase (Fig 3B). After 100 learning episodes, all three agents showed a similar number of steps to reach either of the goals. With regard to pain aversion, the OVaRLAP agent also showed quick learning. The OVaRLAP agent exhibited smaller numbers of wall hits in the early learning phase than did the other agents. However, after 50 learning episodes, the number of wall hits of the OVaRLAP agent was comparable to that of the simple TD learning agent (Fig 3C). By contrast, the MaxPain agent first hit the walls as many times as the simple TD learning agent, but the number of wall hits then quickly decreased. The OVaRLAP agent showed its characteristic performance in the five grid-worlds with different configurations (Fig 3D-3F). These results suggested that the OVaRLAP agent could learn a reward approach and pain aversion in a very efficient manner owing to its generalization capability, whereas it was inferior to the simple TD in long-term reward learning and to MaxPain in long-term pain aversion.

**Fig 3.**
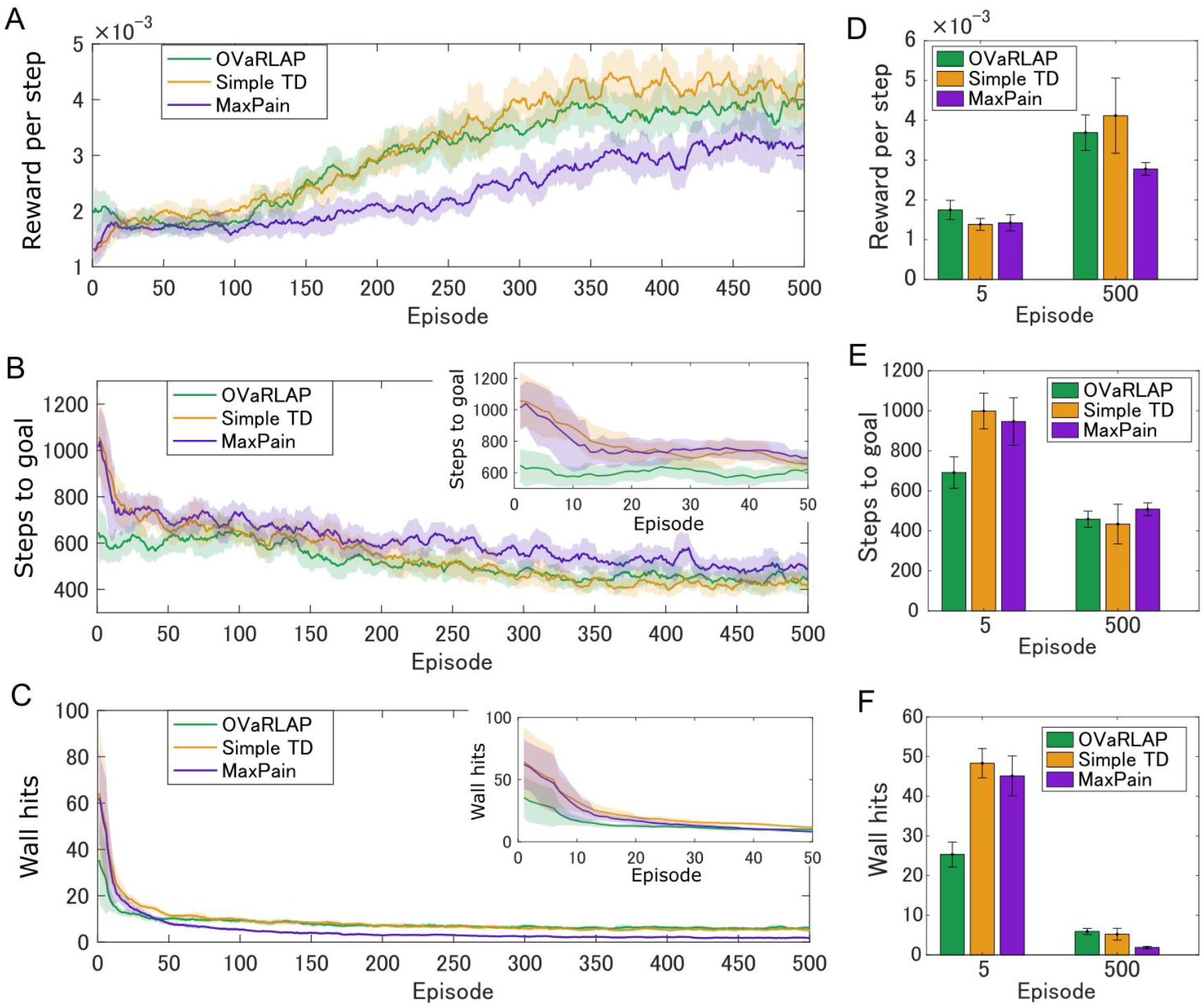
Average performance of OVaRLAP, the simple TD learning, and MaxPain in the painful grid-world navigation task. (**A**) The average reward per step, (**B**) the average number of steps to either of the two goals, and (**C**) the average number of wall hits, based on the three types of agents after each number of learning episodes on the horizontal axis. The grid world shown in Fig 1B was used. The number of steps to either of the two goals and the number of wall hits for the first 50 learning episodes are magnified in the top right insets of panels B and C, respectively. Each average (on the vertical axis) was taken over 50 separate runs and then smoothened as moving averages over nearby 11 episodes (on the horizontal axis). The thick lines represent the moving averages and the shadow areas indicate the moving standard deviations. (**D**–**F**) The average performance over five simulation experiments, each of which used a grid-world with different configurations. The grid world shown in Fig 1B and its variants were used (S1 Fig presents the details of the variants.). (**D**) The average reward per step, (**E**) the average number of steps to either of the two goals, and (**F**) the average number of wall hits, based on the three types of agents after 5 learning episodes and 500 learning episodes, are shown. Each bar indicates the average of five values (i.e., for five different grid-world configurations), each of which is the average over 50 separate runs. Each error bar represents the standard deviation over the five values.

Figure 4 shows how the value function was developed by using the three methods, in which OVaRLAP showed unique profiles (Fig 4, top). First, the values of positions close to the walls decreased, as did the values of the positions of the walls. This generalization constructed a safe passage to the lowly rewarded goal and simultaneously a hazard to approach the risky, but highly rewarded goal, after only five learning episodes. This value hazard completely disappeared after 500 episodes because the reward learning had been propagated to the squares on the passage to the high-reward goal. Thus, quick pain learning was essential for risk aversion and conservative reward-seeking in the early learning stage.

**Fig 4.**
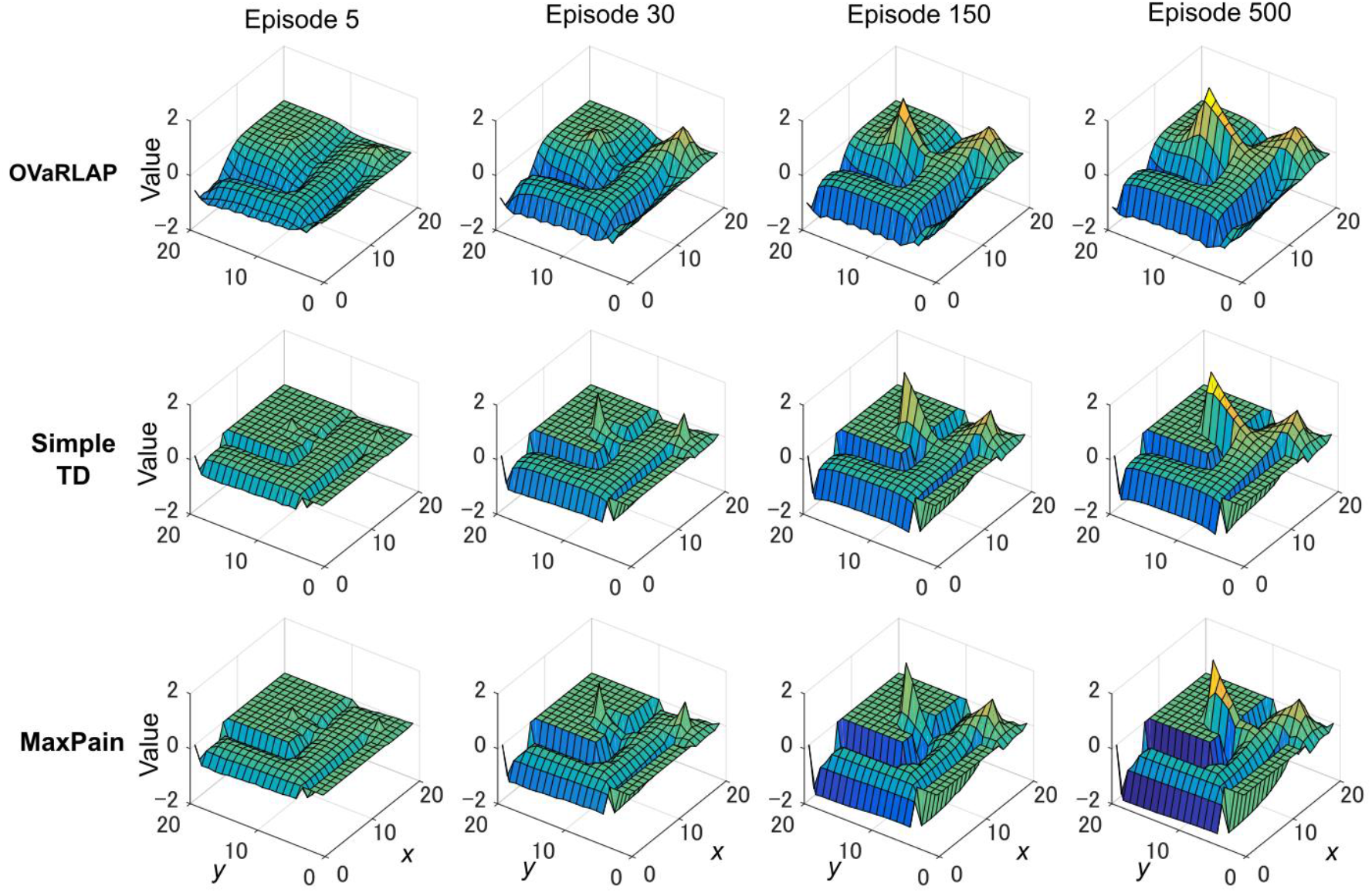
Transition of the value function for each model in the grid-world navigation task. The average state values over 50 separate runs after 5, 30, 150, and 500 learning episodes for OVaRLAP (top row), simple TD learning (middle row), and MaxPain (bottom row), when applied to the painful grid-world navigation task shown in Fig 1B. For MaxPain, the result of the subtraction of the value function for the pain from the value function for the positive reward (i.e., the main factor for decision-making) is displayed.

The simple TD learning algorithm did not construct any value hazard to the high-reward goal during its learning process, because no special system existed for pain learning (Fig 4, middle). The MaxPain algorithm also produced hazards on the passage to the high-reward goal (Fig 4, bottom). In contrast to OVaRLAP, the hazard progressively increased as the learning proceeded and was never flattened owing to its strong ability toward pain learning. For this reason, the MaxPain algorithm persistently maintained low values on the way to the high-reward goal.

### The effect of a disturbed OVaRLAP

We next examined how the learning behaviors of OVaRLAP were disturbed by disabling weight updating by negative TD errors and introducing noise to the fixed network (Fig 1C). The generalization after obtaining a reward became a little noisy because of the noise introduction (Fig 5A). We applied this disturbed OVaRLAP to a grid-world navigation task without pain (Fig 1D). We tested each of the following two-by-two settings: normal or no update for the negative TD error (i.e., “intact” or “impaired”), and with or without noise in the fixed network (i.e., “noised” or “unnoised”).

**Fig 5.**
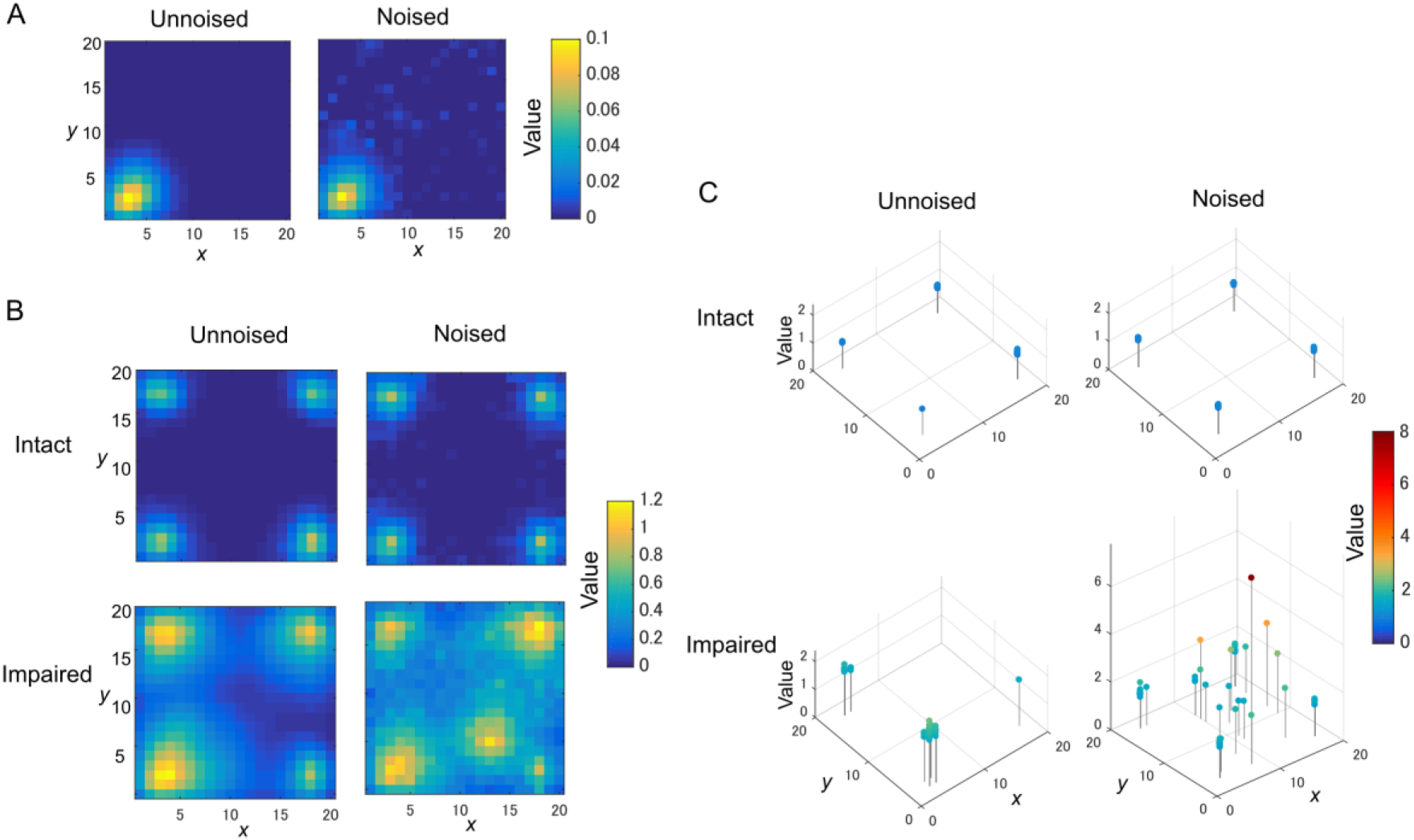
The TD learning task for disturbed OVaRLAP. (**A**) The value function is updated after reaching the left-bottom goal of the grid-world shown in Fig 1D once with OVaRLAP without noise (left) and with noise (right). (**B**) The value function after 120 learning episodes in a single run for OVaRLAP with intact and impaired updates (top and bottom rows, respectively) for the negative TD error with and without noise (right and left columns, respectively) in the fixed connections. (**C**) The maximum value and its position in the value function after a total of 40,000 time steps (378.8 learning episodes on average) in each run. Each panel shows the maximum values in 50 separate runs for OVaRLAP under the following settings: intact or impaired updates (top or bottom row, respectively) for the negative TD error, and without noise or with noise (left or right column, respectively) in the fixed connections. In the noisy case, the noise was independently introduced to the initialization of the fixed connections for each run.

The impaired noised agent acquired an aberrant value function: it increased the value for a specific position distant from the goals, as if the position would give a positive reward (Fig 5B, bottom right). The value function of the intact noised agent was slightly noisy (Fig 5B, top right), but nearly the same as that of the intact unnoised agent (Fig 5B, top left). The impaired unnoised agent made the values around the goals higher than those of the intact agents, but the values for positions distant from the goals remained low (Fig 5B, bottom left).

We ran 50 runs for each setting to confirm the reproducibility. For the noised agent, a different pattern of noise was applied in each run. Fig 5C presents the maximum value and its position in the value function after 40,000 steps in each run. The intact agents always established the maximum values at the positions of goals and maintained them consistent to the actual amount of the rewards (Fig 5C, top row). The maximum values of the impaired unnoised agent were higher than the actual amount of the rewards (Fig 5C, bottom left). Their positions were not necessarily the same as the actual goals, but they were, at most, two steps away from the goals. In contrast to these three agents, the impaired-noised agent yielded maximum values that were quite apart from the actual goals (Fig 5C, bottom right). Their values were higher than the actual amount of rewards and were sometimes exceedingly high.

## Discussion

In this study, we proposed a novel reinforcement learning model named OVaRLAP to analyze how the computational characteristics in the striatal dopamine system affect behaviors. The OVaRLAP model reproduced stimulus generalization and discrimination for reward learning and for punishment learning. If reward learning precedes in a generalized manner, then discrimination is achieved by subsequent value reshaping, based on punishment learning or on the omission of an unpredicted reward. By contrast, if punishment learning precedes and is generalized, then subsequent reward learning or the omission of unexpected punishment causes a discrimination. In the navigation task in the painful grid-worlds (Figs 3 and 4), pain was preceded because it was relatively frequently observed, and the situation was then in the latter case. By contrast, in the painless grid-world (Fig 5), the former case appeared.

The OVaRLAP model enabled safe and efficient exploration in the painful grid-world navigation task. The transition of the value function for the OVaRLAP agent shows how stimulus generalization and discrimination contributed to managing the trade-off between reward-seeking and risk aversion (Fig 4, top).

First, punishment learning due to hitting walls was generalized quickly, as shown in the value function after five episodes. It led to a preference for the center of the passage (i.e., risk aversion). The OVaRLAP agent indeed first showed fewer wall hits than did the simple TD learning and MaxPain agents (Fig 3C and 3F). It contributed to reducing the number of steps to reach either of the goals in the early learning phase (Fig 3B and 3E). The OVaRLAP agent thereafter increased the values of positions close to the walls. This transition indicated that the agent discriminated between the safe and painful positions. This discrimination increased the tendency to reach the high-reward goal. In short, stimulus generalization of punishment learning induced risk aversion, followed by discrimination for switching to reward-seeking. The simple TD learning agent did not show risk aversion (Fig 4, middle). Its value function was optimized to maximize future reward, which is consistent with its high reward per step (Fig 3A and 3D). The MaxPain agent showed strong risk aversion, based on its separate value learning system to expect future pain (Fig 4, bottom row). It achieved very few wall hits (Fig 3B and 3E), but meanwhile, it maintained risk aversion and low values on the way to the high-reward goal, which corresponded to a lower reward per step, compared to the other agents (Fig 3A and 3D).

Stimulus generalization and discrimination were effective in safe and efficient exploration; however, its dysfunction may induce aberrant learning behaviors, as demonstrated by the disturbed OVaRLAP (Fig 5). The impaired update for the negative TD error in combination with the noise in the fixed connections induced aberrant valuation. The value increased by noisy stimulus generalization was not reshaped by discrimination because of the impairment of the punishment learning. A single update of the value was only slightly affected by the noisy stimulus generalization (Fig 5A); however, as the number of arrivals to the actual goals increased, such aberrant updates of the value function would have allotted abnormally high values to some positions that were actually of no reward (Fig 5B-5C).

The structure of OVaRLAP gives insight into the neurobiological basis of stimulus generalization and discrimination. The two pre-output neurons separately responded to positive and negative prediction errors so as to update the connections between the hidden-layer neurons and the corresponding pre-output neuron. This hybrid learning system exhibits good correspondence to the striatal dopamine system in which D1- and D2-SPNs differentially respond to dopamine so that the connections between the cortex and the striatal SPNs are differently modulated [22, 24, 25]. Transforming the input into the hidden-layer activity is an approximation of the process by which an external stimulus and the internal state are encoded into neural activity patterns of the cortex. This process is assumed to be rather independent from dopamine-dependent plasticity. However, this process should not be simply based on random connections because the similarity of the inputs has to be maintained in the encoding process to achieve stimulus generalization. The encoding could instead be learned in a different manner from the reward-related learning. Therefore, the fixed connections between the input and hidden layers in OVaRLAP can be the result of learning for such encoding process. How the brain learns the encoding process is a topic for future research, but some implications are derived from computational models of the primary visual cortex and the primary auditory cortex that adopted unsupervised learning [37, 38].

Compared to ordinary reinforcement learning algorithms, the uniqueness of the OVaRLAP model is that it is characterized by fast generalization and value separation. Its advantage regarding safe and efficient exploration primarily occurred by fast generalization, whereas its potential defect in causing maladaptive behaviors was derived from fast generalization and from value separation. Mikhael et al. [26] previously demonstrated the advantage of value separation. Their model coded reward uncertainty into the sum of the synaptic weights of D1 and D2 SPNs. In addition, their model can adjust the tendency to choose or avoid risky options by changing the weight of the positive and negative values, which is assumed to reflect tonic dopamine level. The further advantages of the value separation were not major topics in our current study, although the OVaRLAP model can also enjoy such advantages. The OVaRLAP model can encode reward uncertainty in distinct connections updated by positive and negative TD errors (red and blue arrows, respectively, in Fig. 1A, left), and can adjust risk-taking tendency by changing the connection weights between the pre-output neurons to the output neuron, which were set constant for simplicity in this study.

The potential defect of OVaRLAP causing maladaptive behaviors can provide a hypothesis on psychiatric disorders. Based on the correspondence between OVaRLAP and the brain, the result for the disturbed OVaRLAP showed the possibility that the noisy encoding process in the cortex and the impairment of dopamine-dependent plasticity induce abnormal stimulus generalization and discrimination, which may underlie delusional symptoms. The relationship between cortical disconnectivity and disrupted learning in schizophrenia has been implied in computational studies [39, 40]; therefore, alterations in cortical connectivity could be a cause of a noisy encoding process. Histologic examinations and diffusion tensor imaging studies for patients with schizophrenia revealed reduced spine density and disrupted white matter connectivity, respectively [41–43]. Reinforcement learning models have been used for attempts to explain positive symptoms, including delusion, in schizophrenia [44–46]. However, these studies do not link abnormal cortical connectivity to positive symptoms. They attribute positive symptoms to aberrant salience (i.e., a surprise response to nonsalient events) [47]. Thus, they focused on an abnormality in the dopamine system, which was also supported by neurobiological evidence [48, 49]. The OVaRLAP model is a new corticostriatal learning model that relates delusional symptoms to an abnormality in cortical connectivity and in the dopamine system.

The relationship between abnormal stimulus generalization and schizophrenia has been investigated by psychological research [3, 32, 33]. The research is based on the hypothesis that abnormal generalization underlies delusional symptoms in schizophrenia. Some studies imply heightened stimulus generalization in schizophrenia. However, the reported abnormality was not sufficiently remarkable for explaining delusion in a straightforward manner. This finding can be attributed to the conditioning paradigm used for their behavioral experiments. Patients with schizophrenia have various deficits in cognitive function; therefore, the experimental design needed to be simple to attribute the result to stimulus generalization rather than other factors. Computational models may potentially be used to investigate how a specific deficit in cognitive function leads to psychotic symptoms through complex learning processes. The OVaRLAP model showed that a small abnormality in stimulus generalization in combination with an unbalanced response to positive and negative prediction error may induce aberrant valuation after reward-related learning, including action selection, state transition, and obtaining rewards from multiple sources.

## Acknowledgement

We would like to thank Hidetoshi Urakubo for his helpful comments.

## Supporting information

**S1 Fig. The grid-world configurations used in the painful grid-world navigation task.** The grid-world shown in Fig 1B is displayed again at the left top. These five grid-world configurations were generated under the following rules, while avoiding symmetric or similar structures. A wide passage consists of rectangles in which the shorter width/height is 5 (based on the number of grids) and the longer width/height is more than 7; the wide passage does not have a dead end. A narrow passage exists with a width of 2 and a length of 6. The starting position and a goal with a lower reward of 1 are located two grids away from the end of the wide passage and three grids away from side walls (i.e., relatively apart from the walls). The passages have to be surrounded by walls. If the passages are separated by walls, the distance between these passages has to be more than three grids. A goal with a higher reward of 2 is one grid away from the end of the narrow passage (i.e., relatively close to the walls). The shortest path to either of the goal requires 26 steps if the agent has to move to the center of the wide passage before turning to the narrow passage.

